# Hand Position Fields of Neurons in the Premotor Cortex of Macaques during Natural Reaching

**DOI:** 10.1101/2024.10.09.617377

**Authors:** Sheng-Hao Cao, Xin-Yong Han, Zhi-Ping Zhao, Jian-Wen Gu, Tian-Zi Jiang, Shan Yu

## Abstract

Planning and execution for goal-directed movements requires the integration of the current position of body or body parts with various kinematic parameters of the movement itself. In the hippocampus, the field-based representation of spatial information plays an essential role in this process during navigation. However, if a similar, field-based encoding framework is also utilized by the motor areas during hand reaching remains unclear. In this study, we investigated the hand position tuning in the dorsal premotor cortex (PMd) neurons (*n* = 601) when four monkeys performing a naturalistic reach-and-grasp task. We show that 132/601 (22%) of PMd neurons increased their firing rates when the monkey’s hand occupied specific positions in the space, forming the “position fields” in their spatial firing maps that can be well described by 2D Gaussian functions/kernels. We further analyzed how the field-tuning of hand position is co-represented with other task-related parameters including the hand moving direction, speed, and reward location in the same population of PMd neurons, revealing a mixed-selective framework also similar to that discovered in the hippocampus. These position-tuned cells demonstrated high efficiency in encoding hand position, with ~10% of overall recorded neurons contributing >80% of accuracy in decoding instantaneous hand moving trajectories. These results suggest that a field-based encoding framework of position may be a common component to representing spatial information and integrating it with kinematic parameters for guiding goal-directed movements of the body or body parts.

## Introduction

An essential function of the brain is to enable the planning and execution of goal-directed movements that can avoid obstacles and approach targets optimally. These movements can occur at various scales, such as the spatial navigation of the whole body and the execution of reaching using the hand. The ability to represent the spatial information of the body ^1–6^ or body parts ^7–10^ and to intergrade it with kinematic parameters is essential for guiding such movements.

For whole-body navigation, this ability is orchestrated by neurons in the hippocampus and its surrounding areas, exhibiting mixed selectivity among various task-relevant parameters, including position^5^, head direction^11^, moving speed^12^, and goal orientation^13^. Specifically, position of the body can be represented by 1) “place cells” in the hippocampus, which discharge when the animal occupies specific locations, giving rise to the concept of “place fields” ^5^, and 2) “grid cells” in the medial entorhinal cortex, characterized by periodic hexagonally spaced place fields^1^. Together, these place fields provide an efficient framework for encoding spatial information, aiding animals in constructing cognitive maps for navigation. For body part movements such as reaching with the hand, it has been known that parameters such as hand moving direction, speed, and goal position are represented through the mixed selectivity of neurons in motor relevant areas^14,15^. Also, neural activities in these areas can be modulated by the spatial location of a reach, either by the starting position^16^, the ending position^17,18^, or a combination of both^19^. However, it remains unclear whether a field-like representation, either place-cell like or grid-cell like, for hand positions exists in somewhere within the motor areas. Resolving this question would offer insights into whether a general representation framework is employed by the brain to guide goal-directed movements across different scales.

To coordinate goal-directed hand movements, a network of various cortical areas is necessary, with the dorsal premotor cortex (PMd) being a crucial node. PMd neurons become active during the delay period before a movement^20,21^, and disruption of PMd activity leads to impaired reaching performance^22,23^. PMd neurons exhibit directional tuning for hand movement^24–27^ similar to that in the primary motor cortex, as well as tuning for other kinematic parameters, including speed^25,28^, distance^29^, and reaction time for a reach^30,31^. Moreover, PMd neurons are influenced by the relative position of the hand, eye, and target, enabling reference frames transformation during planning^32,33^. Given its role in movement planning and the integration of spatial and kinematic information, the PMd presents an appealing target for exploring field-like representation for hand position.

## Results

We performed this study in four male macaques (*macaca mulatta*, Monkey A, B, X, and Z), each of which was implanted with a Utah Array (Blackrock Neurotech) in the left PMd (Figure 1A). The array consisted of 96-channels for monkey X and 48-channels for other monkeys. Neurophysiological activities were recorded as the monkeys performed a natural reach-and-grasp task^34,35^ (Figure 1B), during which they used their right arm to grasp fruit pieces on a wand held by the experimenter. In each trial, the wand was moved to an arbitrarily chosen location, either remaining static or changed to a new position while the monkeys were reaching (see Figure S1 for trajectories of the food). The position of the right hand of the monkeys was reconstructed from video recordings captured by four cameras (Figure S2A). In total we collected 839 (Monkey A, 51; Monkey B, 101; Monkey X, 390; and Monkey Z, 297) putative single units (SUs) during 18 daily behavioral sessions, each session lasting an average of 26.3 ± 6.5 minutes (mean ± SD) and comprising approximately 100 trials. See Table S1 for details of each session.

**Figure 1.**
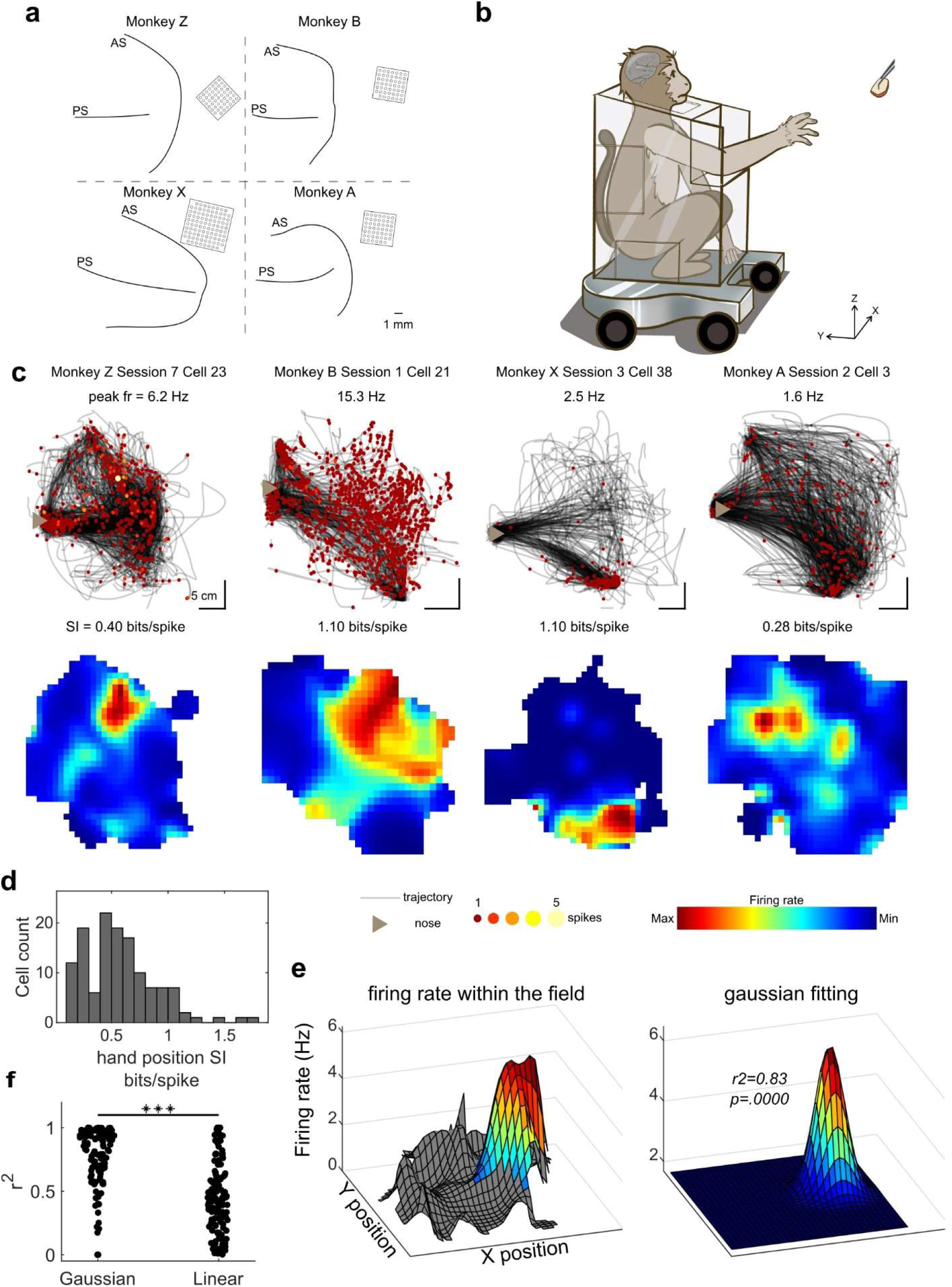
Hand position specific tuning in the dorsal premotor cortex. (**A**) Recording regions mainly covered the hand area of the dorsal premotor cortex in each monkey. Black squares denote position of inserted Utah arrays (96 channels for Monkey X, 48 channels for Monkey A, B, and Z). Black lines denote brain sulci used to help locating the implantation site. PS, principal sulcus; AS, accurate sulcus. (**B**) Diagram of the natural reach-and-grasp task. (**C**) Upper row: trajectories (black lines) of the hand position from four example sessions (one for each monkey). Spikes from four representative cells in each corresponding session are plotted as circular markers. The color and size of the markers are set according to the spike counts. The brown triangle indicates the position of the monkey’s nose (facing right). Peak firing rate for each cell is labeled on the top of panels. The vertical scale bars represent 5 cm along the x axis and the horizontal scale bars represent 5 cm along the y axis in each panel. Bottom row: corresponding 2D spatial firing rate maps, denoted with SI. Dark blue indicates the minimal firing rate within the map and dark red indicates the maximal firing rate within the map. (**D**) The distribution of SI of all hand position-tuned cells (*n* = 132). (**E**) Fitting the hand position firing fields using a 2D Gaussian function. Left, color-coded cell activity of the first cell in (**C**), shown as a function of hand position. Area outside the firing field is masked with grey. Right, fitted gaussian function of the firing field, *r^2^* = 0.83, *p* = .0000. More details of the fitting analysis are described in Methods: Field detection. (**F**) Comparison between the coefficient of determination (*r^2^*) of gaussian fitting and that of linear fitting, *n* = 132, *** *P* < 0.001, paired t-test.

Figure 1C shows trajectories of the hand movements (black lines, projected to the XY plane, i.e., horizontal plane) for an exemplar session from each monkey, superimposed with the spikes of a representative cell. The observed spikes were not random; rather, neurons tended to discharge when the hands were within specific regions. To account for the effect of different time-occupancy in individual positions, we generated 2D-spatial firing rate maps by dividing the spike-count map by the time-occupancy map (Figure S3).

To quantify the spatial specificity of the firings of PMd neurons, we calculated the spatial information (SI)^36^, defined as information content of individual spikes regarding the position of the hand, for each neuron. A hand position-tuned cell was identified if its SI exceeded the 99^th^ percentile threshold of the shuffled data. Figure 1C shows the 2D-spatial firing rate map of four exemplar cells (one for each monkey) that met such criterion. In total, we analyzed 601 (Monkey A, 40; Monkey B, 83; Monkey X, 248; and Monkey Z, 230) putative single units (SUs) with stable spatial firing rate maps (Figure S4). Among them, 132 neurons (22.0%; Monkey A, 4; Monkey B, 34; Monkey X, 57; and Monkey Z, 37) were identified as hand position-tuned cells. This proportion was significantly higher than expected by chance (*P* < 1e-323, Binomial test with expected chance level *P_0_* = 0.01), and the average SI was 0.59 ± 0.29 bits/spike (see Figure 1D for the distribution of SI). These results demonstrate that a sizeable proportion of PMd neurons exhibit significant hand-position specificity in their firing activities. Figure S5 showed the spatial firing rate maps for all hand position-tuned cells.

To examine the field-based tuning of hand position quantitatively, we first identified the hotspots within the spatial firing rate maps (above 50% of the peak firing rates) as the “position field”, similar to the place fields in hippocampus cells^37^. Then we fitted the firing rate *r* within the position field of each cell using a 2D Gaussian function of hand position: 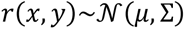 (Figure 1E). As a result, the firing activities of 83% (109 of 132) of the hand position-tuned cells could be well described by a Gaussian function (*r^2^* = 0.80 ± 0.19, *P* < 0.001, F-test). For comparison, we also fitted the firing activities as a linear function of hand position to explore whether these cells encoded spatial gradients at certain orientations, as previously suggested for neurons in the primary motor cortex^38^. Only 50% (66 of 132) of the hand position-tuned cells possessed a planar neuronal response surface within the 2D space (*r^2^* = 0.52 ± 0.24, *P* < 0.001, F-test). More importantly, the coefficient of determination (*r^2^*) of the linear function was significantly lower than that of the Gaussian function (*P* < 1e-25, paired t-test, Figure 1F). The superiority of Gaussian fitting remained in the case of fitting full map data (*P* < 1e-25, paired t-test, Figure S6A). These findings suggest that hand position is indeed represented by a field-like tuning of cells in the PMd. Additionally, we examined whether these position fields exhibited periodic hexagonal spacing and found few grid-like cells (Figure S12).

During the hand reaching movement, in addition to the hand position, other task variables could also affect neural activity, complicating the analysis of spatial firing rate maps. To address this issue, we first calculated the mean vector length, speed modulation depth and food location SI to investigate the selectivity of PMd neurons for hand moving direction^24–27^, speed^25,28^, and food location^39,40^, respectively (see Figure 2A for exemplar cells for each cell type). The population summary of cells with various tuning properties is shown in Figure 2B. We found that PMd neurons exhibited mixed selectivity for diverse task-relevant variables. We identified 24% of PMd neurons (144 of 601) as hand direction-tuned cells and 32% of PMd neurons (192 of 601) as hand speed-tuned cells. We also found 3% of PMd neurons (18 of 601) showed spatial specific firings for food location. Notably, 36% of hand position-tuned cells (47 of 132) showed no significant correlation with other variables.

**Figure 2.**
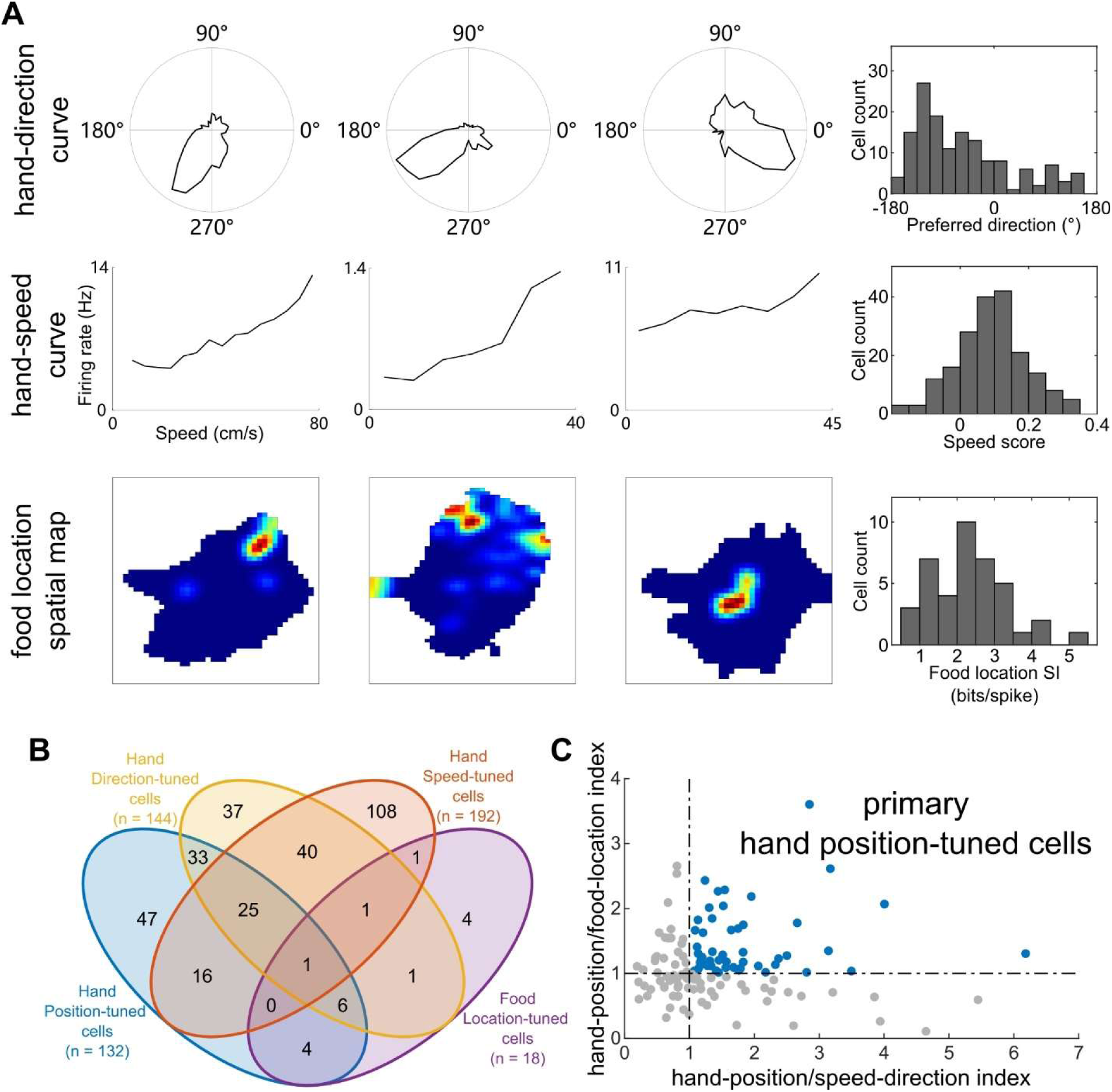
Hand position specific firing in PMd cell is not a byproduct of known tuning properties. (**A**) Firing rate as a function of hand-direction, hand-speed, and food-location in three representation cells respectively. The upper row, firing rate as a function of hand moving direction, followed by the distribution of preferred direction of hand direction tuned cells (*n* = 144). The middle row, firing rate as a function of hand moving speed, followed by the distribution of speed score of hand speed tuned cells (*n* = 192). The bottom row, color-coded spatial rate maps for food location, followed by the distribution of SI of all food location-tuned cells (*n* = 18). (**B**) Population summary of cells tuned to hand position, hand moving direction, hand moving speed or food location. (**C**) Scatter plot showing hand-position/speed-direction indexes and hand-position/food-location indexes of 132 putative hand position-tuned cells met the SI criterion. During the reconstruction analysis, we obtained two normalized reconstruction errors based on two opposing hypotheses and subsequently, the ratio of these errors yields the hand-position index. We identified cells with higher hand-position index (> 1) as primary hand position-tuned cells whose hand position tunings could not be explained by pure other tunings (*n* = 50, blue dots). The vertical dotted line and the horizontal dotted line indicate the identity hand-position index.

Next, to ensure that the spatial specific firings we observed could not be explained solely by the tunings for the direction, speed or food location, we screened hand position-tuned cells through reconstruction analysis^13^ (Figure S7). We conducted a two-step analysis to successively assess the effects of kinematics and food location tunings. First, we retained 80 out of 132 candidate hand position-tuned cells which had lower reconstruction errors assuming pure position tuning (hand-position/speed-direction index > 1, cf. the x-axis in Figure 2C), which suggested that their position tuning property cannot be attributed to the observed speed-direction tuning. Second, out of the remaining 80 cells, we identified 50 (Monkey A, 0; Monkey B, 18; Monkey X, 17; and Monkey Z, 15) cells which had stronger hand position tunings than food location tunings (hand-position/food-location index > 1, cf. the y-axis in Figure 2C), which are referred as primary hand position-tuned cells hereafter. It is noteworthy that we do not suggest exclusive tuning for hand position in these cells. Instead, the designation underscores the hand position tuning as a non-spurious feature within the mixed selectivity framework in the PMd.

After excluding cells predominantly affected by other variables, we reevaluated the fitting results for the spatial tuning function of firing activities. The firing rate of these primary hand position-tuned cells continued to be better described as a Gaussian function of hand position across the space (*r^2^* = 0.78 ± 0.21) rather than a simple linear plane (*r^2^* = 0.54 ± 0.24; *P* < 1e-10, paired t-test, Figure S6B).

Next, we examined other properties of spatial firing rate maps of the primary hand position-tuned cells. The spatial coherence, which measures the smoothness of firing rate maps, was 0.39 ± 0.16 (Figure 3A). The mean spatial sparsity, defined as the fraction of motion space in which a cell is active, was 0.52 ± 0.17 (Figure 3B). The mean area of positional firing fields was 98.7 ± 85.3 cm^2^ (Figure 3C). Multiple fields were observed in 12% (6 of 50) overall significant hand position-tuned cells (Figure 3D). The shapes of fields were predominantly elongated, as indicated by the ratio of the fields’ principal axis lengths being significantly larger than 1 (Figure 3E, *P* < 1e-11, paired t-test). For each monkey, we gathered fields for all primary hand position-tuned cells across sessions and drew them together (Figure 3F), showing that these fields tended to be widely distributed across the reachable space. By summarizing the locations of the fields in the recording sites from all monkeys, there seems to be a tendency for the hand position preferences represented by hand position-tuned cells to correlate with their locations within the cortex: cells located in the rostral part of the PMd predominantly represented the hand positions on the left side of the body, whereas cells closer to ventral premotor cortex and the primary motor cortex showed more preferences for hand positions in front of the body or on the right side (Figure S9).

**Figure 3.**
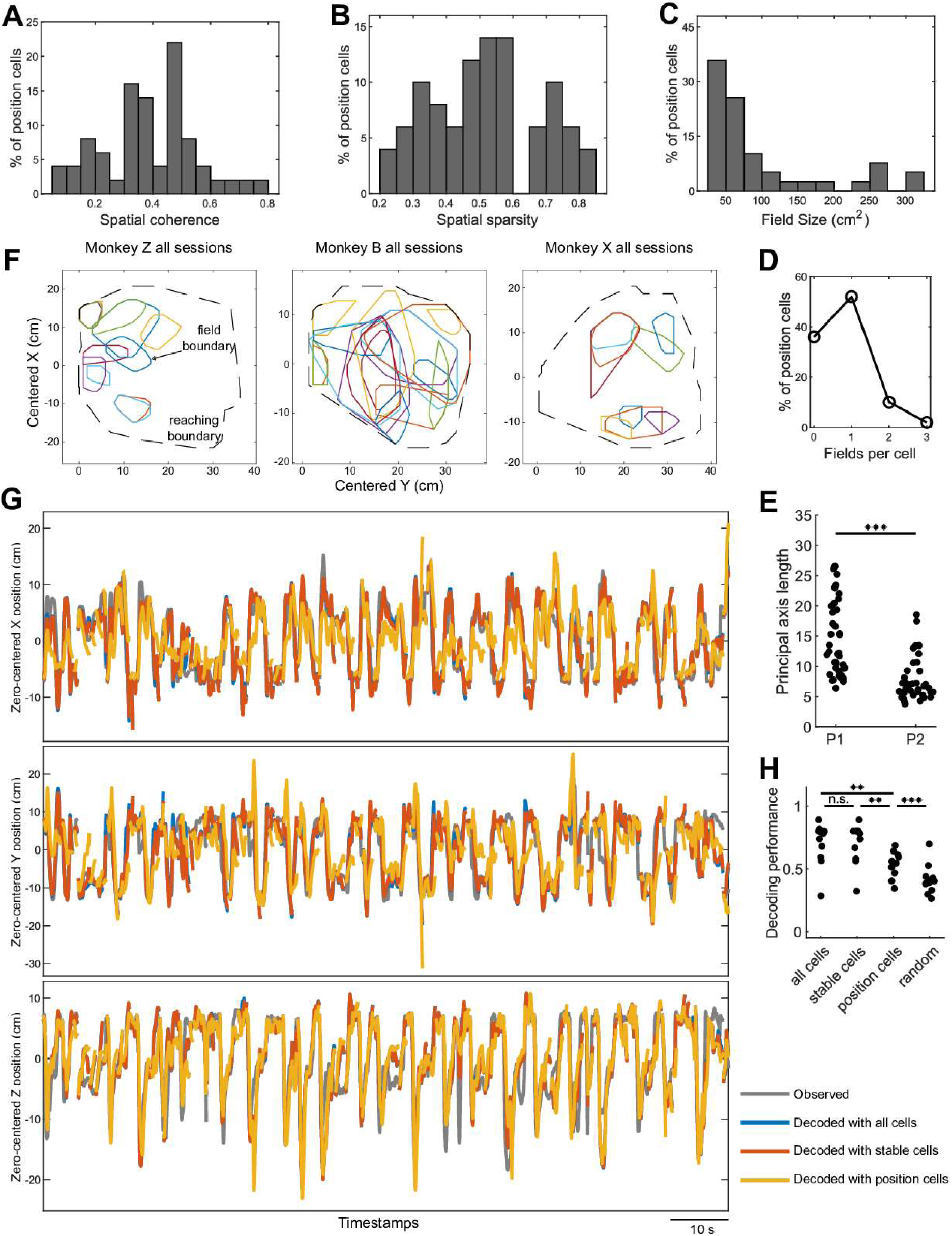
Properties of significant hand position-tuned cells. (**A-D**) Distributions of spatial coherence (**A**), spatial sparsity (**B**), field size (**C**) and number of position fields per cell (**D**) across all significant hand position-tuned cells (*n* = 50). (**E**) Comparison of lengths of two principal axes of position fields. (**F**) Distributions of the position fields across all sessions for each monkey. Colors distinguish fields from different hand position-tuned cells. These positions are relative to the nose of the monkey, which was set as (x, y) = (0, 0). (**G**) Examples of observed and decoded hand trajectories using different cell groups. The time series of positions along the X and Y axis are shown in the upper and middle panel, respectively. Gray lines are the observed trajectories, blue lines are the decoded trajectories using all recorded cells, red lines are the decoded trajectories using cells with stable XY-spatial firing rate maps, and yellow lines are the decoded trajectories using significant hand position-tuned cells. The results shown are based on the entire testing dataset of one session, with the stationary period excluded. The horizontal scale bar represents 10 s. The third panel shows trajectories along the Z axis and the results are calculated from the XZ plane. See Methods: Trajectory decoding analysis for more details. (**H**) Distributions of decoding qualities (Pearson correlation) using different groups of cells and the distribution of the decoding chance performances. The group of all cells--all putative SUs, cell number *n* = 839, averaged 59.2 ± 31.9 across individual sessions; the group of stable cells--cell number *n* = 601, averaged 42.8 ± 18.6 across individual sessions; the group of position cell--significant hand position-tuned cells after reconstruction analysis, cell number *n* = 50, averaged 4.2 ± 2.9 across individual sessions. n.s., not significant, ** *P* < 0.01, *** *P* < 0.001, paired t-test.

To further examine how the population of primary hand position-tuned cells could represent the hand positions across the reachable space, we used these cells to decode the hand moving trajectory (cell number *n* = 50, averaged 4.2 ± 2.9 across individual sessions, see Figure 3G for the observed and decoded trajectories). Overall, the correlation coefficient (*cc*) between the observed and decoded trajectories across all sessions in all animals is 0.54 ± 0.10 (Figure 3H). We also evaluated the decoding performances using the group of all putative SUs (*cc* = 0.68 ± 0.17, cell number *n* = 839, averaged 59.2 ± 31.9 across individual sessions) or the group of cells with stable spatial firing rate maps (*cc* = 0.68 ± 0.16, cell number *n* = 601, averaged 42.8 ± 18.6 across individual sessions). Although lower than the performance using all cells (*P* < 0.01, paired t-test) or stable cells (*P* < 0.01, paired t-test), the primary hand position-tuned cells, which amount to only 10% of the recorded neurons, achieved 84% of the decoding performance obtained by using all cells and 82% of the decoding performance obtained by using all stable cells. These results suggest that the primary hand position-tuned cells represent the spatial information during reaching highly efficiently.

## Discussion

Graziano et al. observed that long-period electrical microstimulation of various sites in the precentral cortex (including the premotor and the primary motor cortices) induced the movement of the monkeys’ hands to specific spatial positions^41^. Our identification of field-based tuning for both hand and target positions reveals the spatial information encoding framework that likely underlies the observed effects of microstimulation. The field-based tuning of hand position we found in the premotor area appears different compared to that observed in the primary motor area and parietal cortex, which has been suggested to exhibit gradient tuning, i.e., cells encode not specific hand positions but rather position gradients along certain orientations^38,42,43^. We note that the reaching task employed in our study encompasses a considerably broader reachable space compared to that utilized in previous works and aligns with the natural conditions of monkey’s arm movements better. In a confined space, the field-based tuning can appear as the linear gradient tuning along certain orientations. Consequently, future research efforts may be warranted to reassess the hand position tuning in the motor area under more natural and expansive conditions.

Several studies have shown that the spatial positions of a reach movement in the 3D space would modulate existing tuning properties in motor areas, such as direction tuning^19,27,44^, distance tuning^16^. Supposing 3D spatial information is similarly encoded in neural activities in the form of position field, discretely sampling trajectories on the 3D spatial firing rate maps would provide a possible explanation for such observations.

As previous studies have revealed the important role of the PMd in coordinates transformation^32,33^, the current finding of the hand position representation in the PMd provides a straightforward way to realize such transformation computationally. By integrating the spatial information of target/stimuli in the PMd and related areas^45,46^ and the hand position represented in this study, it is possible to obtain a vector representation of the direction and distance of a reaching movement through making populational subtraction based on the same spatial coordinate. This subtraction operation achieves a transformation from body-centered (position coordinates of the hand and the target) to limb-centered (direction and distance relative to the hand) reference framework. Then downstream cortical areas could further translate such information into intrinsic motor commands. Importantly, to provide stable information for the movement of a body part, we argue that the field-like hand position coding in the PMd is egocentric and would remain constant after body relocation, which is distinct from the allocentric space representation for body navigation.

The simplified, natural reaching tasks we employed lacked explicit and distinguishable periods demanding for preparations or plannings. This leaves the questions of how these hand position tuned cells are involved into the broader functions because PMd plays a significant role in many motor cognitive processes including preparation and execution of movement^47–49^, abstraction^50^, decision signals^51,52^ and motor learning^53^ etc. We derived our main findings based on the non-stationary period data, another decoding analysis (Figure S8B) showed that populational hand position coding remained stable and would not be affected by the hand motion state, suggesting that the representation of current hand position by these cells was not dependent upon the presence of motion itself. But it awaits future studies to reveal how the position coding framework we found here may contribute to the state estimation or motor planning, for example, exploring the dynamic properties^54–57^ of hand position tuning, in which the coding is calculated for not only the instantaneous hand position but also the future and past positions.

Elucidating the specific tuning function of individual neurons can yield more insights into representational geometry and neurocomputing^58^. For example, it has been demonstrated that the nonlinear conjunctive selectivity of multiple features ensures improved reliability and efficiency in information representation compared to the pure selectivity^59^. In this context, the extrinsic space can be decomposed into two separate dimensions (or three dimensions in the 3D space). As a result, the theory of mixed-selectivity suggests that field-based tuning of position, which is a nonlinear mixed representation of various space dimensions, provides computational advantages over position gradient tuning, which entails pure selectivity along a single dimension. The functional benefits of field-based tuning may account for the observed similarities in the tuning for 1) hand position during limb movements, 2) body position during navigation, and 3) the relative position of various concepts in an abstract cognitive map^60,61^. The advantages of field-based tuning in representing information are consistent with the high efficiency demonstrated in decoding hand trajectories in the present study, as well as the high efficiency in representing spatial or abstract conceptual information within a more general framework termed the Tolman-Eichenbaum Machine^63^, suggesting that functional optimization may lead to a similar framework for information representation in different brain regions. In the same light, the mixed selectivity among position and kinematic parameters can be regarded as a more sophisticated conjunctive representation, constituting a higher-level representation framework shared between the hippocampus and motor-related areas to guide goal-directed movements.

## Limitations of the study

There are several limitations of the current study. First, we did not track the eye movements during the reach-and-grasp task. During the planning and execution of reaching movement, ocular position can modulate neural activities in PMd^32^. In the current experiment, the usual pattern we observed was that when the food appeared in sight, monkeys immediately looked at the target, not their hands. When there was no food in sight, the monkeys’ gaze directions were more or less random, and their hands often put somewhere on the primate chair. The lack of correlation between gaze direction and hand position suggested that the tuning of the latter was not very likely be qualitatively affected by eye movements. Nevertheless, eye movements should be considered in future experiments. Second, upper limb movements naturally occurred in the 3D space, but accurate estimation of the firing activities in the 3D space requires significantly more sampling data than in the 2D case. Therefore, in the current paper, we focused only on the 2D plane, to which the conclusions can be made with strong data support. The nature of hand position fields in the 3D space remains to be unveiled.

## STAR★Methods

### Key resources table

| REAGENT or RESOURCE | SOURCE | IDENTIFIER |
| --- | --- | --- |
| Experimental models: Organisms/strains |  |  |
| Rhesus macaques |  | Macaca Mulatta |
| Software and algorithms |  |  |
| MATLAB R2020a or R2022b | Mathworks | https://www.mathworks.com/ |
| Offline Sorter x64 V4 | Plexon | https://plexon.com/products/offline-sorter/ |
| Fieldtrip | Fieldtrip Developers | https://www.fieldtriptoolbox.org/ |
| Deeplabcut | Deeplabcut Developers | https://github.com/DeepLabCut/DeepLabCut |
| Pose3d | Pose3d Developers | https://github.com/SwathiSheshadri/pose3d |
| Venn diagram | Venn diagram Developers | https://www.mathworks.com/matlabcentral/fileexchange/116760-venn-diagram |

### Resource availability

#### Lead contact

Further information and requests for resources should be directed to and will be fulfilled by the lead contact, Shan Yu.

#### Materials availability

Not applicable. This study did not generate reagents.

#### Data and code availability

- All data that support the findings for this study are available from the lead contact upon request.
- Custom code used for analyzing spike activities and motion data are available from the lead contact upon request.
- Any additional information required to reanalyze the data reported in this work paper is available from the lead contact upon request.

#### Experimental model and study participant details

Four male rhesus macaques (Macaca mulatta), aged from 9 to 11 years were used in this study. All animal experimental procedures were performed in accordance with international and national guidelines for the administration of experimental animals under a protocol approved by the Biomedical Research Ethics Review Committee of the Institute of Automation, Chinese Academy of Sciences.

### Method details

#### Behavioral task

Before the start of the experiments, all animals were surgically implanted with a titanium head post. Monkeys were trained to sit head-fixed in a primate chair for behavioral testing and electrophysiological recording.

The behavioral task was a natural reach-and-grasp task^34,35^. Specifically, fruit pieces were attached to one end of a wand manually held by the experimenter. The monkeys were allowed to use their right arm to reach and grasp the fruit pieces and then feed themselves. There was no explicit instruction or time limit on performing the task, which allowed the animals to choose how to reach-and-grasp at their own pace once the fruit pieces were presented. The wand was moved to random locations in front of the monkey, either staying static or changing to a new position during the reaching. The food might be moved outside the reachable range of monkeys to encourage their maximum reach movements. Importantly, these periods constituted only a very small proportion of each session as monkeys would not reach out if they found the food beyond their reaching limits for too long. There were no explicit constraints, except for those imposed by sitting in the primate chair with head fixed (Supplementary Video 1), on the spatial range of reaching for monkeys during the task. The recorded spatial ranges in the 3D space for each monkey were: Monkey A, 35cm × 37cm × 42cm; Monkey B, 35cm × 35cm × 39cm; Monkey X, 39cm × 38cm × 50cm; and Monkey Z, 44cm × 38cm × 50cm. In each daily session, about ~100 pieces of fruits were given, which lasted for 9.5 ~ 35.7 mins. Each monkey was repeatedly tested for 3-8 sessions.

#### Electrodes and implantation surgery

Intracortical microelectrode arrays (Utah Array, inter-electrode distance: 400 µm, electrode length: 1 mm, Blackrock Neurotech) targeting the hand area of the left dorsal premotor cortex (PMd) were implanted using standard stereotaxic procedures under isoflurane anesthesia. Animals were fasted for 24 hours before the surgery. Ketamine (10 mg/kg) was given intramuscularly for initial sedation and anesthesia. Then, animals were transferred to a dedicated surgery room and fixed in a stereotaxic apparatus. Gas anesthesia was performed with oxygen (30% partial pressure), nitrous oxide (70% partial pressure), and 1%-2.5% isoflurane through endotracheal intubation. After the craniotomy, the principal sulcus and the arcuate sulcus were identified. A multi-channel array (96-channels for Monkey X, 48-channels for Monkey A, B, and Z) was then implanted in the PMd. Postoperative routine antibiotic therapy continued for 5-7 days. All animals had a recovery period of 1-2 weeks before the recording sessions.

#### Hand motion tracking

Motions of the right hand of the animals were recorded using four cameras (Hikvision) from different angles (Figure S2A). The video data were sampled at 60-100 frames/sec. We firstly used markerless tracking software *DeepLabCut* (DLC)^62^ to extract the 2D positions of the right hand (the metacarpophalangeal joint of the middle finger) and the food in each video frame and then reconstructed the 3D positions out of high confidence 2D data points (likelihood ≥ 0.9, DLC estimated a likelihood value for each data extracted) using the toolbox *pose3d*^64^. In addition, we extracted the positions of monkeys’ noses referred to as the position of the monkeys. Position data were then post-processed to fill small gaps (< 150 ms) by median filtering. Camera calibration parameters for each camera pair were estimated after each recording session via the *Computer Vision Toolbox* from Matlab (Mathworks Inc.). We excluded the stationary periods during which the speed was lower than 1 cm/s to avoid the effects of changes in moving status^12^.

#### Neural recording and spike sorting

Neural data were recorded during the task. We used a CerePlex Direct system (sample rate: 30 kHz, Blackrock Neurotech) or an Intan Recording Controller (sample rate: 20 kHz, Intan Inc.) to save the raw, broad-band signals recorded from the implanted electrodes. Units were extracted offline by the automatic spike-sorting software—Mountainsort^65^. A fifth-order Butterworth bandpass (250-7500 Hz) filter was applied to the raw signal, followed by a zero-phase component analysis (ZCA) whitening across channels. The detection threshold was set to −5.5 s.d. and the waveform length was 1.6 ms (−0.4 ms ~1.2 ms). We further verified the sorted units manually in Offline Sorter (Plexon Inc.) and identified putative single units according to two criteria: a) 2-ms inter-spike interval (ISI) violation rate < 2%; b) a clear clustering and separation of waveforms could be seen in at least one feature space. Only single units with an average firing rate > 0.1 Hz were included for further analyses.

Given that the recordings were obtained from chronically implanted electrode arrays, we also considered a case where unique cells were recorded repeatedly over multiple sessions and explored the effect on the hand position tuning. Please see Supplementary Information and Figure S10-11.

#### Synchronization between videos and neural recordings

The camera (Hikvision) we used supports hardware-trigger mode, that is, when the camera receives a rising edge voltage (TTL 0 to TTL 1), it takes one frame image. We used a signal generator (DG1022U, RIGOL Technologies) to generate a periodic square wave signal to control the frame rate of the camera. We connected four cameras to the same output port of the signal generator, so we could record multiple videos from different angles simultaneously. Similarly, the trigger-in port of the neural recording system (CerePlex Direct or Intan Recording Controller) was also connected to the same output port of the signal generator, as a result, when a rising edge was generated, cameras captured one frame image while the neural recording system recorded the timestamp of this rising edge which also referred as the timestamp of this frame along with neural data.

#### Calculating motion data and firing rate

After extracting the position data from the videos, we could further calculate the moving velocities and moving directions in each frame easily, so we got the position train, speed train and direction train. We used the triggered timestamps in neural data to set the time bins to count the number of spikes resulting in the corresponding spike count train. We could divide the spike count train by the frame rate of videos to get the spike firing rate. We used the position trains and corresponding spike count trains to visualize the spike-trajectory plots and calculated the spatial firing-rate maps further. We used the speed trains and firing rate trains to calculate the speed-tuning curve. We used the direction trains and firing rate trains to calculate the mean vector length. We used the position trains and firing rate trains in our trajectory decoding analysis.

#### Spatial firing-rate map

Analyses were conducted in the XY (i.e. horizontal) plane as most reaching movements were performed along this plane.

##### Hand

Spatial firing-rate maps were constructed for each putative single unit. Specifically, the spike counts and the time occupancy of the right hand in each spatial bin (1 cm × 1 cm) were calculated, generating one spike-count map and one time-occupancy map. Each map was then smoothed by a 2D Gaussian filter (standard deviation: 1.5 bin, Matlab function *imgaussfilt*). We then computed the firing-rate maps used for further analyses by dividing the spike-count map by the time-occupancy map (Figure 1D). To ensure enough sampling time, invalid spatial bins with time occupancy less than 50 ms were excluded from the 2D firing-rate maps.

##### Food

We calculated the reward firing rate map for each neuron as a function of the food location. We excluded data after the monkey grasped the fruit in each trial because the position of the monkey’s hand and food overlapped during this period. Then the computing process was the same as above.

#### Recording stability

To test the stability of spatial firing rate maps, we divided all recorded data within each session into two equal halves. Two new firing-rate maps were computed respectively with the same process mentioned above. The Pearson correlation, *r*, between the two maps was calculated. Cells with low stability, i.e. *r* < 0.3, were excluded from further analyses^66^ (Figure S4).

#### Field detection

Firing rate maps were binarized by thresholding at 50% of the peak firing rate. Then we defined the firing field as a set of contiguous regions whose area was larger than 25 cm^2^ and was visited by the hand at least five times^37^. Field properties such as boundary, size, principal axis length, and eccentricity were obtained by using Matlab function *regionprops* applied to the binary maps. *Fitting the activity function within the firing field.* Assum that the firing rate *r* is a 2D gaussian function of the hand position 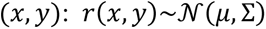.

First, we rotate the positional coordinate axis by an angle of *θ* to eliminate the correlation between them:

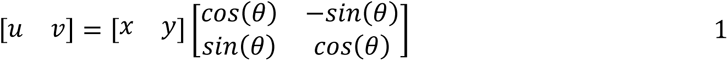

Then, we fitted the firing rate *r* as a function of (*u*, *v*) based on the equation below:

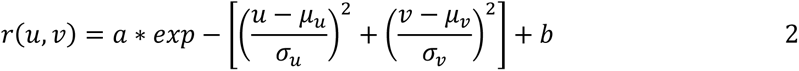

where *a, b* are gain and baseline, *μ*_u_/*μ*_v_ is the mean value along each axis and *σ*_u_/*σ*_v_ is the standard deviation along each axis.

We extracted the rotation angle *θ* by using Matlab function *regionprops*. The fitted gaussian function was obtained by Matlab function *fit*. As a comparison, we directly used Matlab function *regress* to calculate the linear regression results for the firing fields. For spatial firing rate maps with multiple fields, we only conducted regression analyses on the main firing field which included the peak firing value.

#### Spatial sparsity

Spatial sparsity was used to reflect the fraction of the motion space in which a cell is active^67^, which was calculated as follows:

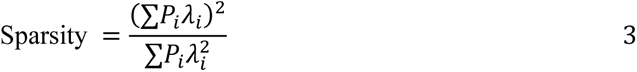

where *P*_i_ denotes the occupancy probability in the *i*th bin, *λ*_i_ denotes the firing rate of the cell in the *i*th bin.

#### Spatial coherence

Spatial coherence was estimated as the mean correlation between the firing rate of each bin and the mean firing rate in the eight adjacent bins^68^, which was used to measure the similarity of local firing rates to that of the neighboring bins.

#### Measurements used for cell type classification

*Spatial mutual information.* The information theory method was used to quantify the amount of information transmitted by the firing of a neuron (bits/spike) regarding the interested experimental variables^2,36,69^. Specifically, we calculated the spatial mutual information as:

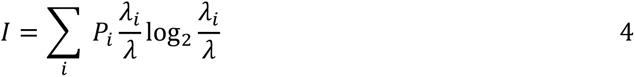

where *P*_i_ denotes the occupancy probability in the *i*th bin, *λ*_i_ denotes the firing rate of the cell in the *i*th bin and *λ* denotes the overall mean firing rate of the cell.

SIs were calculated based on the hand spatial firing rate maps to select putative hand position-tuned cells and were also calculated based on the food spatial firing rate maps to identify the food location-tuned cells. We employed a bootstrap procedure^70^ to verify that the estimation error arising from finite sampling is negligible in the current datasets (data not shown).

##### Speed modulation depth (to identify hand speed-tuned cells)^12^

We first calculated the speed-tuning curve according to the time series of moving velocity and the firing rate. The equally spaced bin size is 5cm/s. The modulation depth of a cell was defined as the difference between the maximum and minimum firing rates in the speed-tuning curve. We also computed the speed score as the Pearson correlation between the time series of the velocity and the firing rate.

##### Mean vector length (to identify hand direction-tuned cells)^12^

First, the circular hand direction tuning curves (angle bin width π/16, Gaussian smoothing, standard deviation: 1.5 bin, Matlab function *smoothdata*) were calculated. Then, the mean vector length was computed as follows:

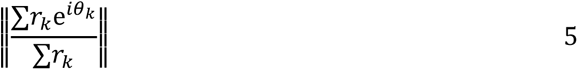

where *θ*_k_ is the angle expressed in radians in the *k*th bin and *r*_k_ is the corresponding firing rate. ||*|| indicates the modulus.

#### Shuffling and classification criterion

To obtain the shuffled data, spike trains were shifted by a random interval that was more than 30 s and less than the session duration minus 30 s (wrapping the end circularly to the beginning). Then we re-calculated all measures mentioned above and repeated this procedure 500 times. If the raw measure was larger than the 99^th^ percentile of the shuffled data, the cell was classified as the corresponding cell types and the chance level was set as *P_0_* = 0.01. Measurement comparison was done within each session. It is worth noting that the nature of the speed modulation depth does not allow for the mixture of information coming from different cells, so that every individual cell had its own threshold.

#### Firing rate map reconstruction analysis

Inhomogeneous reaching behavior could mislead our judgment of hand position-tuned cells. For example, if a cell is only tuned for hand moving direction and speed, and the monkey’s hand always passes through certain spatial locations with specific speed and/or direction, then an apparent tuning for spatial locations could be seen. To exclude such possibility, the reconstruction analysis^11,13^ was adopted to examine how well the position tuning we observed can be explained by the kinetic parameter tuning properties. Specifically, this analysis was carried out by a two-step process: 1) assuming that the cell is only jointly tuned to speed-direction and reconstructing the spatial firing rate map based on this assumption; 2) assuming that the cell is only tuned to hand location and reconstructing the speed-direction firing rate map. The reconstructed spatial firing rate map was computed using the following equation (Figure S7):

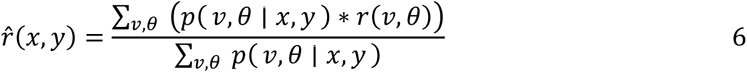

where *p*(*ν*, *θ* | *x*, *y*) is the fraction of time spent at a specific speed-direction bin (*ν*, *θ*) while the hand in a particular spatial bin (*x*, *y*), and *r*(*ν*, *θ*) is the firing rate for this speed-direction bin.

The reconstructed speed-direction firing rate map was computed using the following equation:

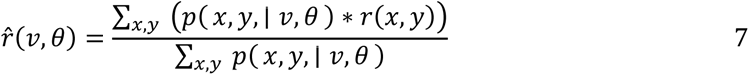

where *p*(*x*, *y*, | *ν*, *θ*) is the fraction of time spent at a specific spatial bin (*x*, *y*) while the hand in a particular speed-direction bin (*ν*, *θ*), and *r*(*x*, *y*) is the firing rate for this spatial bin.

The reconstruction error was defined as the normalized mean squared error (Eq. 8) between the observed tuning map and its reconstructed version. All maps were normalized by their maximal firing rate.

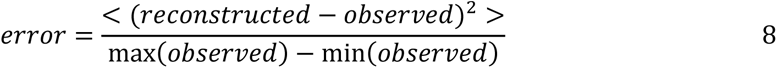

We compared the errors of the two reconstructions mentioned above. In other word, we calculated the *hand-position/speed-direction index* = *error under speed-direction-tuning assumption* / *error under hand-position-tuning assumption*. The reconstruction with the lower error compared to its counterpart indicates that the property used to perform this reconstruction explains the activity tuning better than the other way around, for example hand-position index greater than 1 for stronger hand position-tuned cells.

We adopted the same reconstruction analysis utilizing hand spatial firing rate maps and reward spatial firing rate maps to assess the interference from food location tunings.

##### Speed-direction firing rate map

We computed speed-direction firing rate maps to represent joint speed-direction tuning properties. The spike counts and time occupancy in each equally spaced bin (bin size: 5 cm/s for speed and π/8 for direction) were calculated. Each map was smoothed by a 2D Gaussian filter (standard deviation 1.5 bin). We then calculated the firing rate maps by dividing the smoothed spike-count map by the smoothed time-occupancy map. Bins in which time occupancy was less than 50 ms were excluded from further analyses.

#### Trajectory decoding analysis

A linear Kalman filter was used to predict the trajectories of the monkey’s hand during each session. Firing rates were calculated in partially overlapping 100 ms bins successively shifted by 50 ms (we downsampled the frame rate of motion data to 20 Hz for convenience). The *state* vector of the Kalman filter *X*_t_ = [*pos vel acc*]^T^ describes the kinematic parameters of the hand in each time step *t*, and the *observation* vector of the Kalman filter *Z*_t_ = [*z*_l_ *z*_2_ … *z*_c_]^T^ represents the firing rates of *c* selected cells.

We used three different groups of neurons to decode the trajectories in each session: group 1-all cells, all the putative single units after spike sorting; group 2-stable cells, cells with stable hand spatial firing rate maps; group 3-primary hand position-tuned cells, cells met the identification criteria. Details of modeling and parameter estimation can be found in a previous study^71^. We utilized 70% of the data starting from the session beginning to train the model and the remaining 30% of the data for testing. Different Kalman filters were constructed for each 2D plane. The averaged correlation coefficients between the predicted and observed trajectories in test datasets were used to quantify the decoding performance.

We directly applied the trained decoders mentioned above to the data collected during stationary periods (defined as movement speed < 1 cm/s) to assess their performance during these stationary periods.

##### Chance level

To get the decoding chance performances, we performed the same decoding process using randomly selected neurons. For example, suppose we have identified *n* primary hand position-tuned cells within a specific session. Subsequently, we randomly selected *n* cells from the pool of stable cells recorded from the same session and evaluated their trajectory decoding performance. This process was repeated for 100 times for this session and the averaging performance was calculated as the final chance performance.

## Acknowledgments

We thank Dong-Lin Gu, Peng-Chen Pan, Kai-Xi Tian, Cheng-Teng Jiang, and Jing-Wen Guo for technical assistance during the experiments. We thank Wen Ren for help with setting up the camera systems. We appreciate professional care provided to the animals by Yan-Yan Liu and Bao-Jiang Niu. We thank Rui Chen for drawing the diagram of the behavioral task. We thank Hong-Dian Yang, Cheng-Lin Miao, Da-Jun Xing, and He Cui for their comments on the manuscript. This work was supported in part by the STI 2030—Major Project (2021ZD0200402, 2021ZD0200200), the International Partnership Program of the Chinese Academy of Sciences (CAS) (173211KYSB20200021), and the Strategic Priority Research Program of CAS (XDB32040200).

The graph abstract was created with BioRender.com.

## Author contributions

S.C. and S.Y. conceived and designed the study. S.C., X.H., and Z.Z. performed the implantation surgeries. S.C. and X.H. performed the behavioral experiments and data recording. S.C. and Z.Z. preprocessed the neural recordings. S.C. wrote the analysis code, analyzed data, and visualized results. All authors interpreted and discussed results. S.C. and S.Y. wrote the manuscript with contributions from all coauthors. All authors reviewed and commented the manuscript.

## Declaration of interests

The authors declare no competing interests.

## Declaration of generative AI and AI-assisted technologies in the writing process

During the preparation of this work the authors used ChatGPT in order to improve language and readability. After using this tool/service, the authors reviewed and edited the content as needed and take full responsibility for the content of the publication.

## Notes

### Competing Interest Statement

The authors have declared no competing interest.

