## Supplementary Information for "Hand Position Fields of Neurons in the Premotor Cortex of Macaques during Natural Reaching"

### Stability of hand position coding across states

Because movements were towards moving targets, it would be intriguing to assess the effect of the dynamic movement planning, such as inhibitory control[1], on the hand position coding in premotor cortex. We chose to explore the stability of coding under different states of food/target.

The state of food was not constant during single trial, so there were no perfect “static-target trials” or “moving-target trials”. Instead, we define the time periods when the food moving speed was smaller than 1 cm/s as the “target static periods” and the remaining time periods as the “target moving periods”. The target static periods accounted for only 2.2±1.6% of each recording session, making it difficult to calculate a valid hand position spatial firing rate map, as in the main text. As a result, we could not determine the stability of hand position coding by directly comparing spatial firing rate maps from two periods. Instead, we decided to utilize a trajectory decoding test:

First, we divided the target moving periods into the training set (70%) and the testing set (30%). Second, we trained a hand trajectory decoder (Kalman Filter, details in Methods) with the training set data and calculated the decoding performance in the test set. Third, we directly applied the same trained decoder to the target static periods to assess the performance. Finally, we adopted the same chance level for decoding analysis in the main text.

The results (Figure S8A) showed that decoders extracted from the target moving periods performed even greater in the target static periods, which revealed the stability of the hand position coding.

Similarly, we divided each session into “hand moving periods” and “hand static periods” and then performed the same analysis process above and arrived at a similar conclusion (Figure S8B).

These two results together proved that populational hand position coding during the reach-and-grasp task was stable and was not affected by the food/target state or the hand state.

**Stability of hand position coding across time**

Specific to the Utah array, we adopted Fraser et al. 's method[2], which integrated pairwise cross-correlogram, autocorrelogram, waveform shape, and mean firing rate as identifying features of a neuron. When comparing two units recorded on different sessions using these features, their similarity scores typically fell into one of two categories: high, suggesting that the recordings came from the same unit, or low, indicating that they originated from different units.

Out of the 839 putative single units extracted in our experiments, this method identified 451 (Monkey A-30, B-54, X-238, Z-129) unique neurons. Further, 53.4% (241/451) of these unique neurons were recorded on only one session and 46.6% (210/451) were recorded repeatedly over multiple sessions.

Out of the 451 unique cells, we selected 340 cells with stable firing rate maps and identified 98 hand position tuned cells following the same process in the main text. Based on this result, the proportion of hand position tuned cells in the PMd came rise to 28.5%. We presented the firing rate maps of these hand position-tuned cells repeatedly recorded in multiple sessions together in Figure S10.

We further explored whether the properties of these hand position tuned cells would change over time. To quantify the stability of the hand position tuned cells, we calculated the correlation between maps of the same unit recorded in different sessions. Also, we calculated the distance between the peak value position of the maps of the same unit recorded in different sessions to measure the drift of this unit's position preference over time. For comparison, we randomly sampled pairs of different units and then computed the above two metrics and sampled 100 such pairs for each monkey. Before the calculations, maps from different sessions were aligned according to the reference point (the location of the monkey).

Comparisons showed that the identified unique units had significantly stable hand spatial firing rate maps across sessions (*P* < 0.001, Kruskal-Wallis test) and significantly smaller drift in position preferences (*P* < 0.001, Kruskal-Wallis test, Figure S11). It is important to note that we did not use hand spatial firing rate maps and peak value positions as features when tracking units across sessions. The results indicated the stability of hand position-tuned cells across sessions.

**Searching for grid-like cells with periodic hexagonal spaced fields in PMd**

To assess whether the hexagonal firing patterns characteristic of grid cells were present in the PMd cortex, we calculated grid scores based on rotated autocorrelograms (see *Grid Score* below). Only one cell had a grid score above the $99^{th}$ percentile of shuffled grid scores (within the horizontal plane), and this percentage was not significantly higher than the chance level (Fig. S12; $p = 0.98$, Binomial test with expected $P_0 = 0.01$). Analyses of the other two planes give the same conclusion. Future investigations should encompass a broader range of sites within the motor control circuit to explore whether such cell type exist.

*Grid Score*. First, we calculated a spatial autocorrelogram based on Pearson's product-moment correlation coefficient with corrections for edge effects and unvisited locations. In practice, the autocorrelogram was estimated as:

$$
\begin{aligned}
S_1 &= \sum\lambda(x,y)\lambda(x-\tau_x, y-\tau_y) \\
S_2 &= \sum\lambda(x,y) \\
S_3 &= \sum\lambda(x-\tau_x, y-\tau_y) \\
S_4 &= \sum\lambda(x,y)^2 \\
S_5 &= \sum\lambda(x-\tau_x, y-\tau_y)^2 \\
r(\tau_x,\tau_y) &= \frac{nS_1 - S_2S_3}{\sqrt{nS_4 - S_2^2}\sqrt{nS_5 - S_3^2}}
\end{aligned}
\tag{1}
$$

where $\lambda(x,y)$ is the average firing rate at position (*x, y*), $\tau_x$ and $\tau_y$ are spatial lags and $n$ is the number of pixels in $\lambda(x,y)$ for which firing rate estimated for both $\lambda(x,y)$ and $\lambda(x-\tau_x, y-\tau_y)$. Autocorrelations were not calculated for lags ($\tau_x$, $\tau_y$) where $n < 20$.

The gird score was based on calculating the Pearson correlations between the autocorrelogram and its rotated version [3], which was a modified version of methods used in [4–7]. First, a series of rings centered on the central field in the autocorrelogram were determined, which has the same inner radius $r_i$ and various out radius $r_o$. $r_i$ was defined as either the first local minimum in a curve showing correlation as a function of the average distance from the center or as the first incidence where the correlation was negative, whichever happened first. $r_o$ was increasing from a minimum of 4 bins more than the inner radius to a maximum of 4 bins less than half of the behavioral border length. The Pearson correlations between the original ring and its rotated versions were calculated. In one group, the rings were rotated for 60 and 120 degrees, and in the other group, the rings were rotated for 30, 90, and 150 degrees. For each ring, the minimum difference, $\tau$, between any of the correlation coefficients in the first group and that in the second group was saved. Then the grid score was defined as the largest $\tau$ among all rings.

**Table S1. Detailed experimental information.** Subjects, task durations, frame rates for motion tracking, and the number of putative single units of each experiment session.

| Subject | Date | Duration | Frame rate | Sorted units |
|---|---|---|---|---|
| Monkey A | 2022.01.12 | 25m8s | 60 | 16 |
| | 2022.01.13 | 27m46s | | 16 |
| | 2022.01.14 | 21m34s | | 19 |
| Monkey B | 2021.07.07 | 34m17s | 100 | 37 |
| | 2021.07.08 | 35m42s | | 22 |
| | 2021.07.09 | 31m49s | | 42 |
| Monkey X | 2021.06.23 | 22m21s | 75 | 81 |
| | 2021.07.01 | 9m27s | 100 | 89 |
| | 2021.07.07 | 34m20s | | 90 |
| | 2021.07.08 | 29m33s | | 130 |
| Monkey Z | 2021.10.16 | 21m6s | 100 | 51 |
| | 2021.10.17 | 28m37s | | 53 |
| | 2021.10.18 | 19m47s | | 51 |
| | 2021.10.19 | 21m21s | | 23 |
| | 2021.12.29 | 28m12s | 60 | 35 |
| | 2021.12.30 | 28m11s | | 31 |
| | 2021.12.31 | 31m10s | | 29 |
| | 2022.01.05 | 22m32s | | 24 |

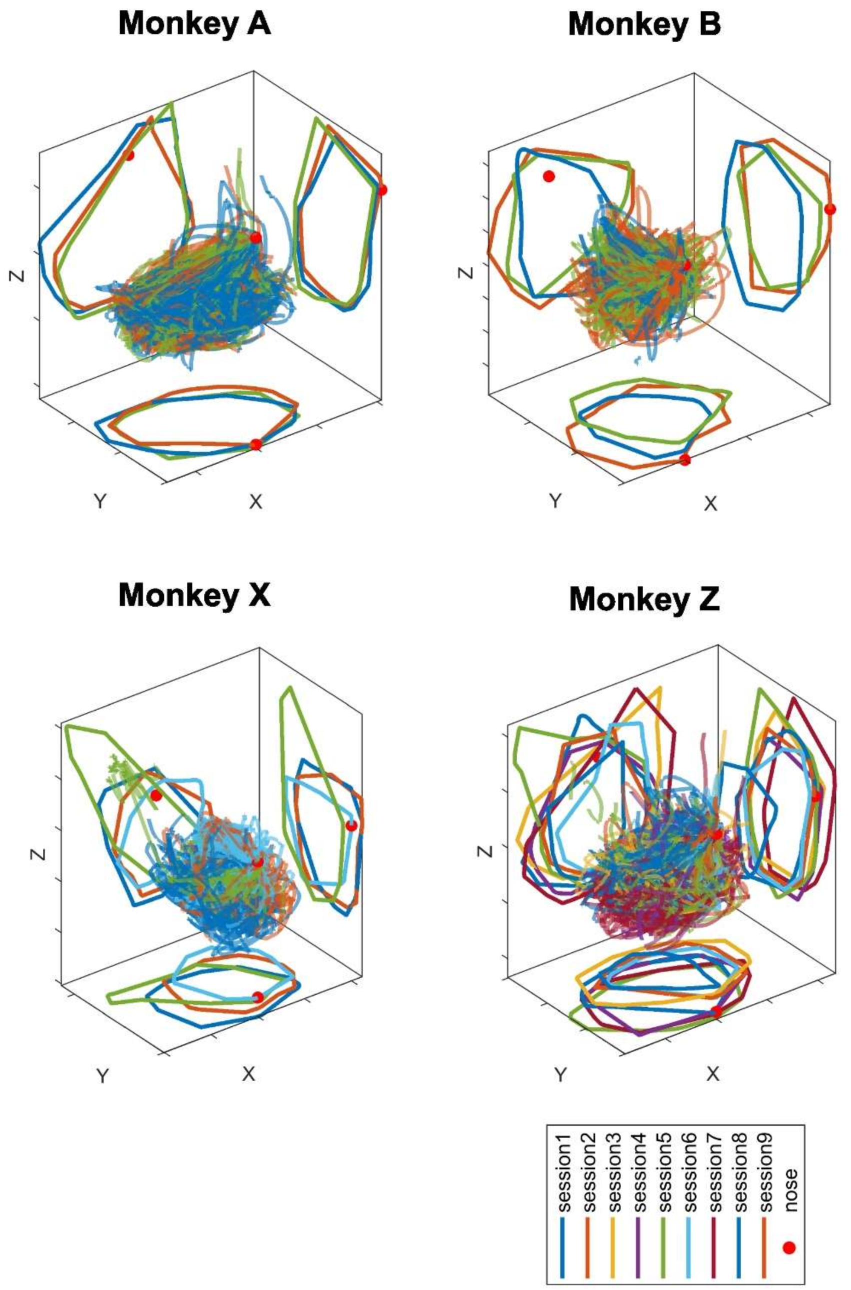


**Figure S1. The spatial trajectories of the reaching targets.** We plot the trajectories of the food with reference to the position of monkey's nose from all sessions into the same panel. Different colors indicate different sessions. The red dot indicates the monkey's nose. Both trajectories in 3D space and boundaries projected into three 2D planes are shown. The experimenter made sure that targets were within the reach of monkeys most of the time.

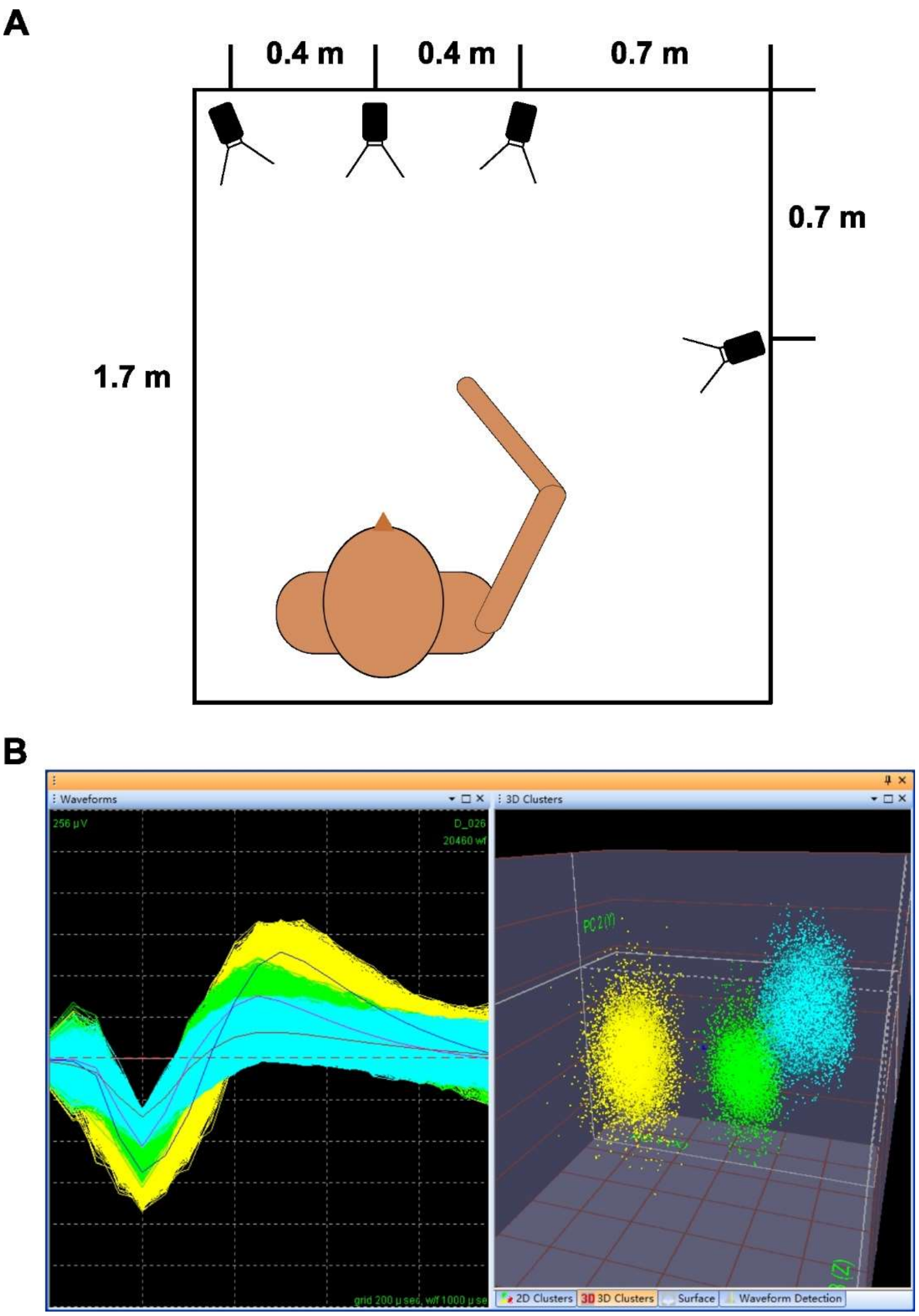


**Figure S2. Camera setting ups and spike sorting criteria.** (**A**) The top view of the motion tracking setup, cameras are ~1 m above the ground. (**B**) Spike sorting illustration for an example single electrode, waveforms and first 3 PC clusters of the three isolated units in this electrode are shown.

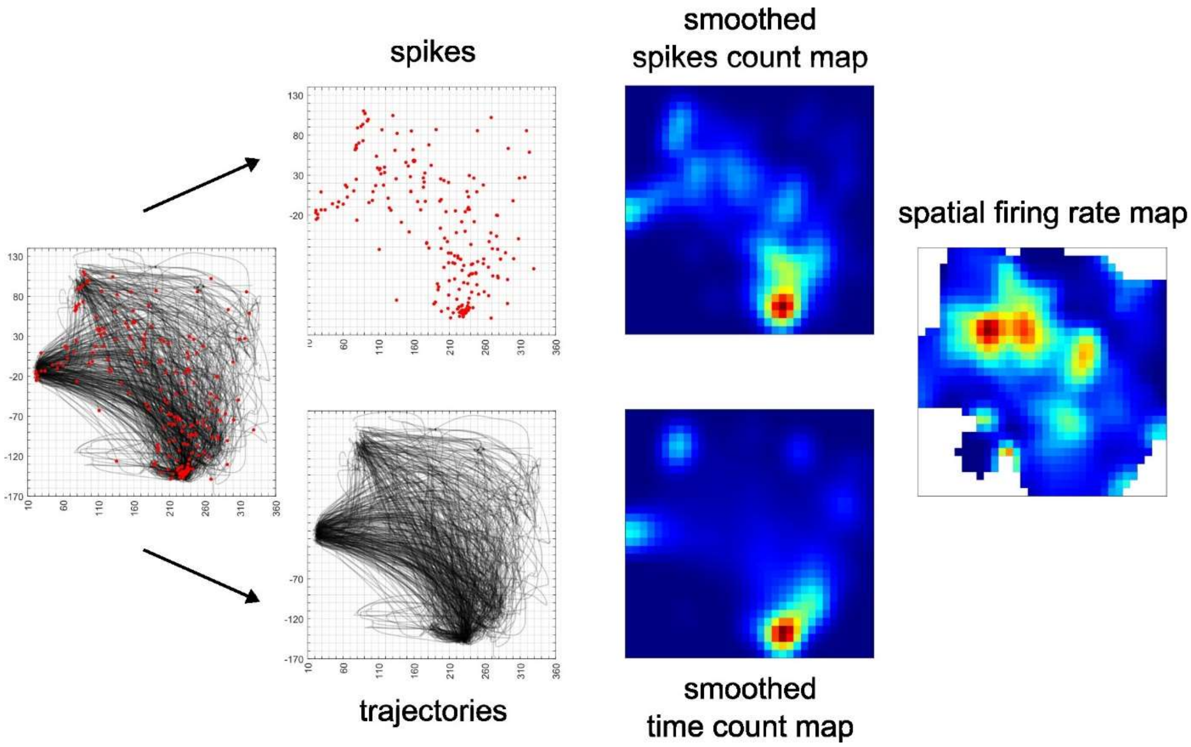


**Figure S3. Illustration of the intermediate steps of the calculating spatial firing rate maps.** Details are described in STAR Methods: Spatial firing-rate map. We first divided the moving space into grided spatial bins (bin size: 1cm × 1cm, column 1 and column 2). Next, we counted the number of spikes and the time in each bin respectively and smoothed the spike count map and the time map using a Gaussian filter (std: 1.5, column 3). Finally, we divided the smoothed spike count map by the smoothed time map to get the spatial firing rate map (column 4).

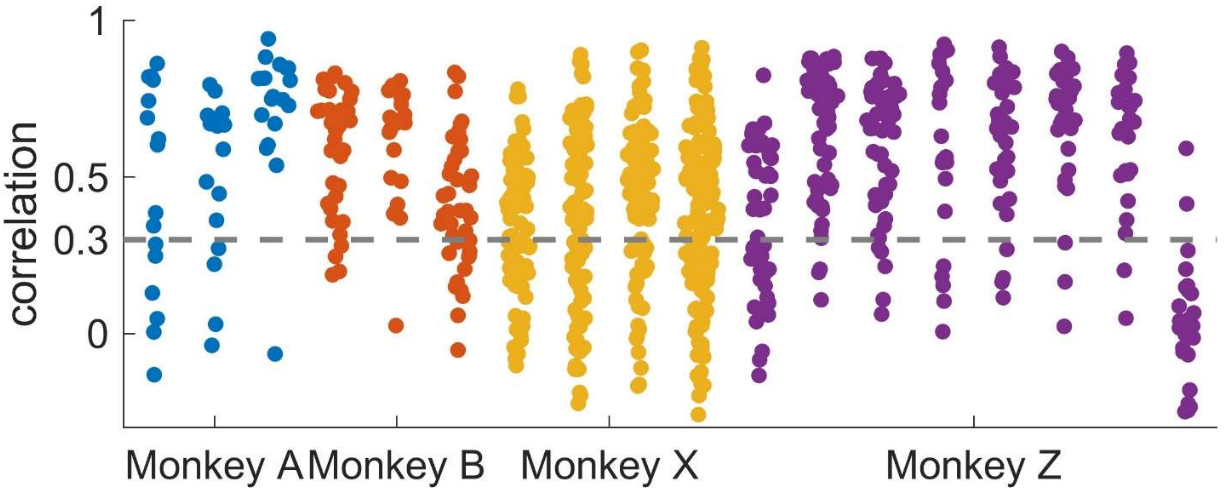


**Figure S4. Distribution of cells' spatial firing rate map stability in each session.** Details of calculating the map correlation was described in Methods. We set 0.3 (dashed line) as the stability threshold. Most sessions of Monkey A (blue dots), B (red dots), and Z (purple dots) showed a bimodal distribution, with 0.3 being a suitable threshold to remove the small peak. The distributions of Monkey X (yellow dots) were unimodal, and we utilized the same threshold to remove neurons with bad recording stabilities only.

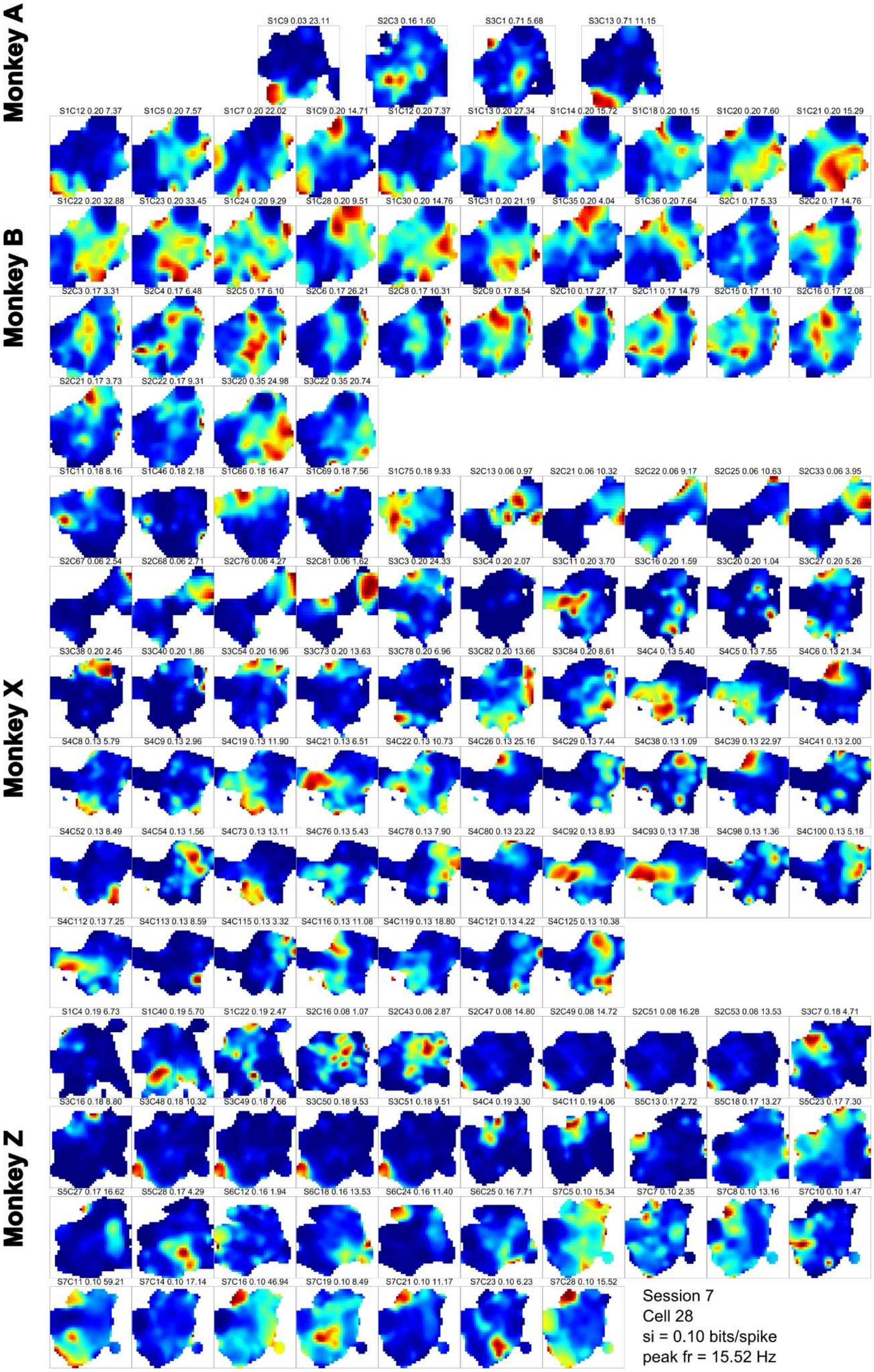
Monkey A
Monkey B
Monkey X
Monkey Z
Session 7
Cell 28
si = 0.10 bits/spike
peak fr = 15.52 Hz

**Figure S5. Spatial firing rate maps of hand position-tuned cells (*n* = 132).** Dark blue indicates the minimal firing rate within each map and dark red indicates the maximal firing rate within each map. Labels, spatial information, and peak firing rate of each cell are marked on the top of maps. The location of the monkey's nose (not actually shown in the figure) is around the left middle of each map, facing right.

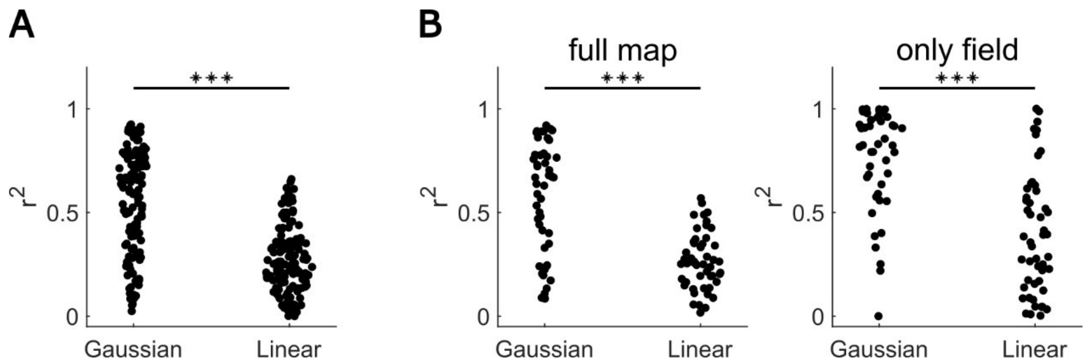


**Figure S6. Comparisons between gaussian fittings and linear fittings.** (**A**) Results in full map data (*n* = 132) still confirmed the superiority of the Gaussian fitting. In the fitting process, only a single Gaussian function was utilized, and a more complex Gaussian Mixture Model could further improve the superiority. (**B**) Comparison of fitting results after reconstruction analysis. The firing rate of primary hand-position-tuned cells (*n* = 50) were better described as a Gaussian function of hand position rather than a liner plane. Left, fitting full map data. Right, fitting position field data. *** $P < 0.001$, paired t-test.

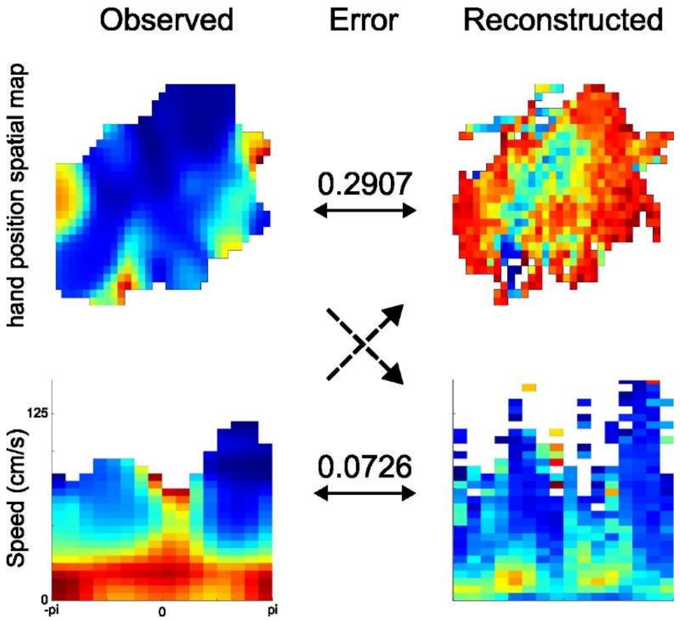


**Figure S7. Illustration of the reconstruction analysis.** Details are described in Methods: Firing rate map reconstruction analysis. Upper row: spatial firing rate maps. Bottom row: speed-direction joint firing rate maps. Left column: observed firing rate maps. Right column: reconstructed firing rate maps. We first assumed that the cell is a pure hand position-tuned cell and based on this assumption and monkey's actual hand moving parameters, we reconstructed what would be the expected speed-direction tuning (upper left to bottom right); and conversely, we assumed that the cell is a pure speed-direction cell and reconstructed the expected hand position tuning (bottom left to upper right). The dashed arrows indicate the direction of reconstruction. Normalized mean squared errors were computed between the observed and reconstructed firing rate maps and indicated between the maps, and the final hand-position/speed-direction index of this cell is 4.0. The calculation of hand-position/food-location index followed a similar process by setting the food-location tuning as the opposing tuning property.

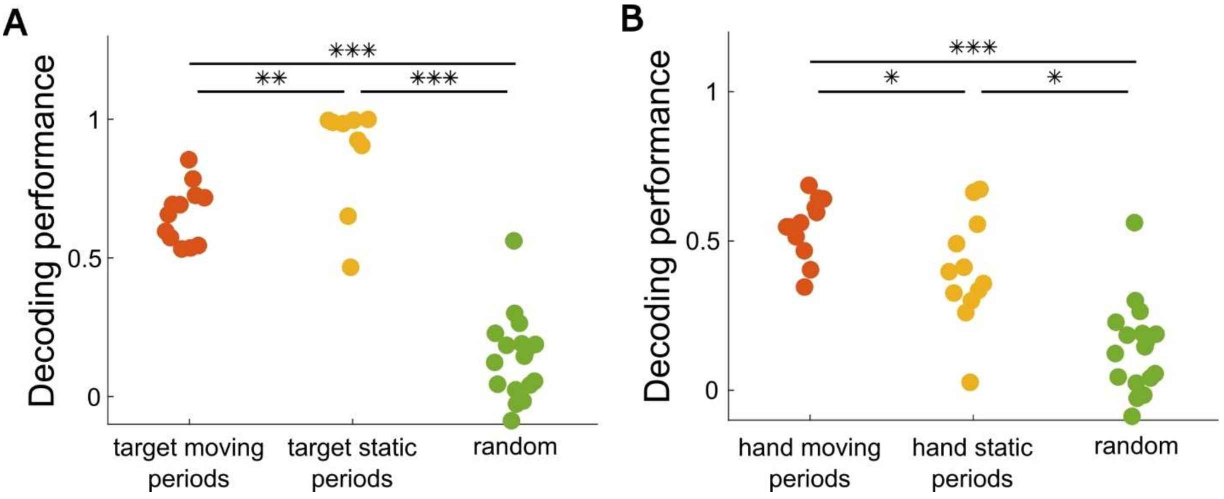


**Figure S8. Comparison of decoding performance across states.** (**A**) The decoding performances in the target static periods (yellow dots) were greater than those in target moving periods (red dots) using the same decoders trained in target moving periods and greater than the random performances (green dots). (**B**) The decoding performances in the hand static periods were lower than those in hand moving periods using the same decoders trained in hand moving periods but greater than the random performances. * $P < 0.05$, ** $P < 0.01$, *** $P < 0.001$, paired t-test.

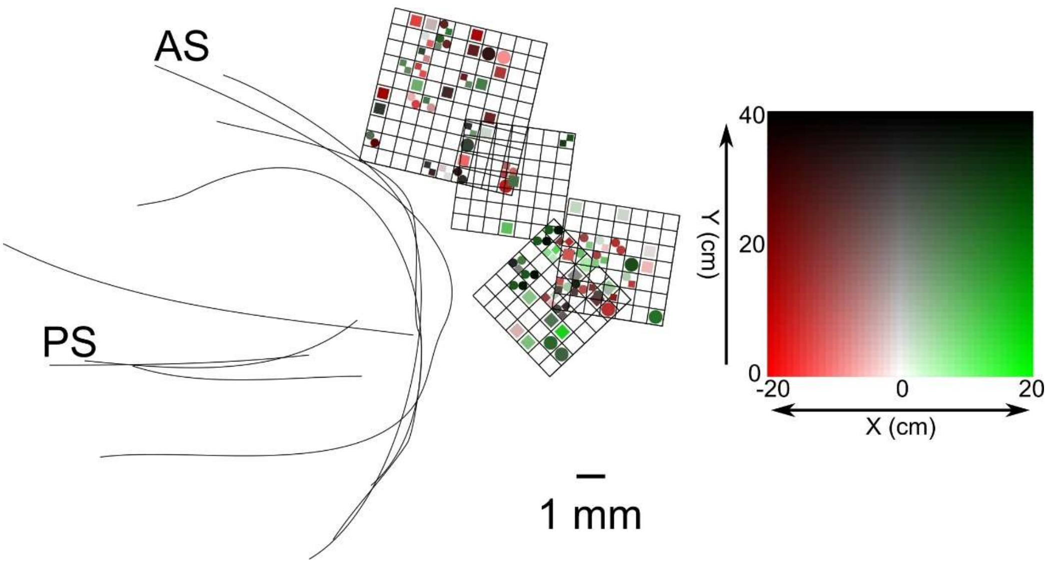


**Figure S9. Hand position preferences in the recording sites summarized from all monkeys and all sessions.** Red denotes the left side of the monkey and green denotes the right side of the monkey. Saturation increases from the midline towards the left/right reaching boundaries, and brightness decreases from near area to far area. Monkeys' body positions were set at (0, 0). Circular markers represented primary hand position-tuned cells (*n* = 50) and square markers represented remaining hand position-tuned cells (*n* = 82). If multiple hand position-tuned cells were recorded at the same electrode site, the size of the markers were adjusted to remain within the same grid.

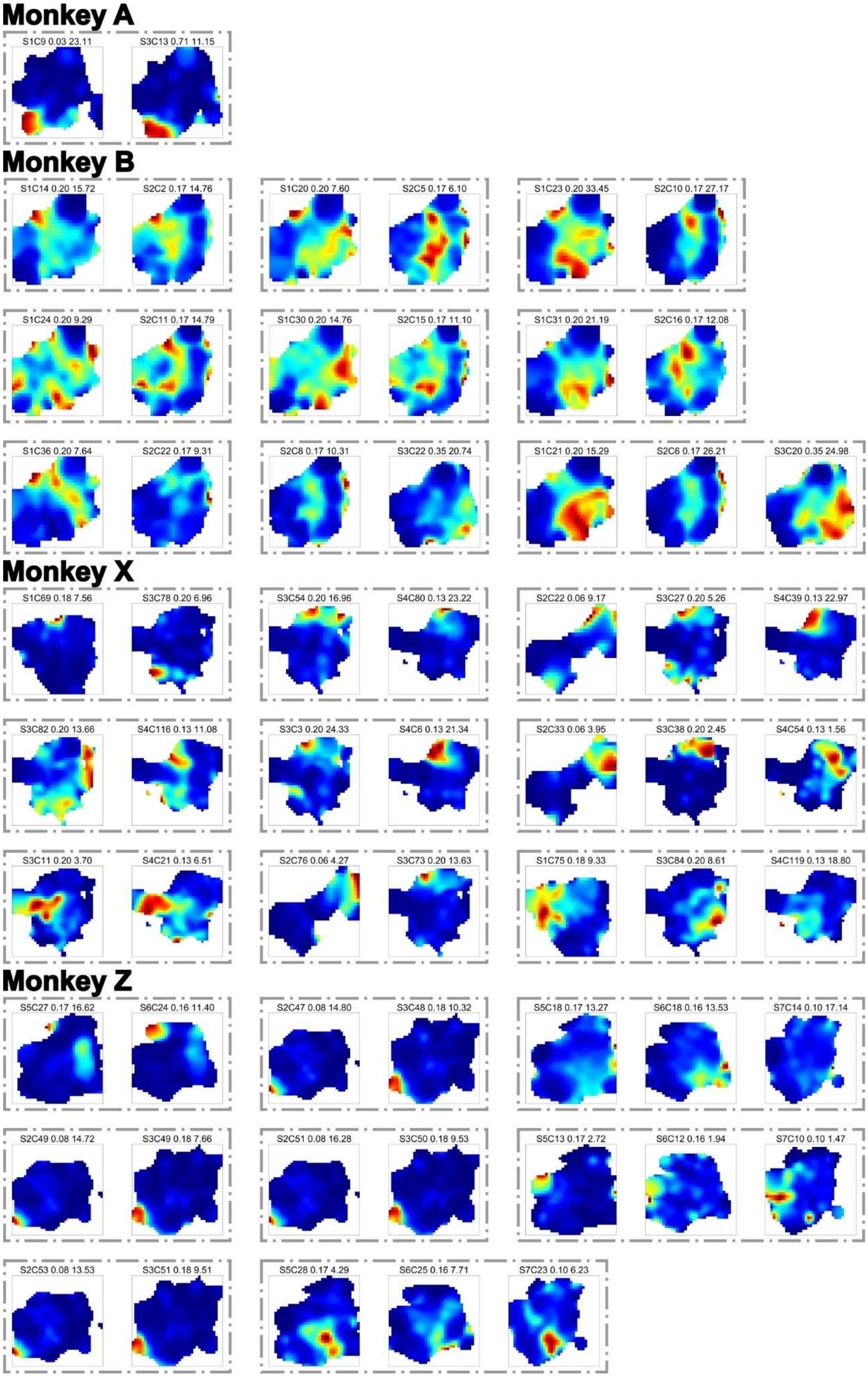
Monkey A
S1C9 0.03 23.11
S3C13 0.71 11.15
Monkey B
S1C14 0.20 15.72
S2C2 0.17 14.76
S1C20 0.20 7.60
S2C5 0.17 6.10
S1C23 0.20 33.45
S2C10 0.17 27.17
S1C24 0.20 9.29
S2C11 0.17 14.79
S1C30 0.20 14.76
S2C15 0.17 11.10
S1C31 0.20 21.19
S2C16 0.17 12.08
S1C36 0.20 7.64
S2C22 0.17 9.31
S2C8 0.17 10.31
S3C22 0.35 20.74
S1C21 0.20 15.29
S2C6 0.17 26.21
S3C20 0.35 24.98
Monkey X
S1C69 0.18 7.56
S3C78 0.20 6.96
S3C54 0.20 16.96
S4C80 0.13 23.22
S2C22 0.06 9.17
S3C27 0.20 5.26
S4C39 0.13 22.97
S3C82 0.20 13.66
S4C116 0.13 11.08
S3C3 0.20 24.33
S4C6 0.13 21.34
S2C33 0.06 3.95
S3C38 0.20 2.45
S4C54 0.13 1.56
S3C11 0.20 3.70
S4C21 0.13 6.51
S2C76 0.06 4.27
S3C73 0.20 13.63
S1C75 0.18 9.33
S3C84 0.20 8.61
S4C119 0.13 18.80
Monkey Z
S5C27 0.17 16.62
S6C24 0.16 11.40
S2C47 0.08 14.80
S3C48 0.18 10.32
S5C18 0.17 13.27
S6C18 0.16 13.53
S7C14 0.10 17.14
S2C49 0.08 14.72
S3C49 0.18 7.66
S2C51 0.08 16.28
S3C50 0.18 9.53
S5C13 0.17 2.72
S6C12 0.16 1.94
S7C10 0.10 1.47
S2C53 0.08 13.53
S3C51 0.18 9.51
S5C28 0.17 4.29
S6C25 0.16 7.71
S7C23 0.10 6.23

**Figure S10.** Rearrangement of spatial firing rate maps of hand position-tuned cells which were identified as the same unit across sessions ($n$ = 27). Legends of the maps are the same as in Figure S5.

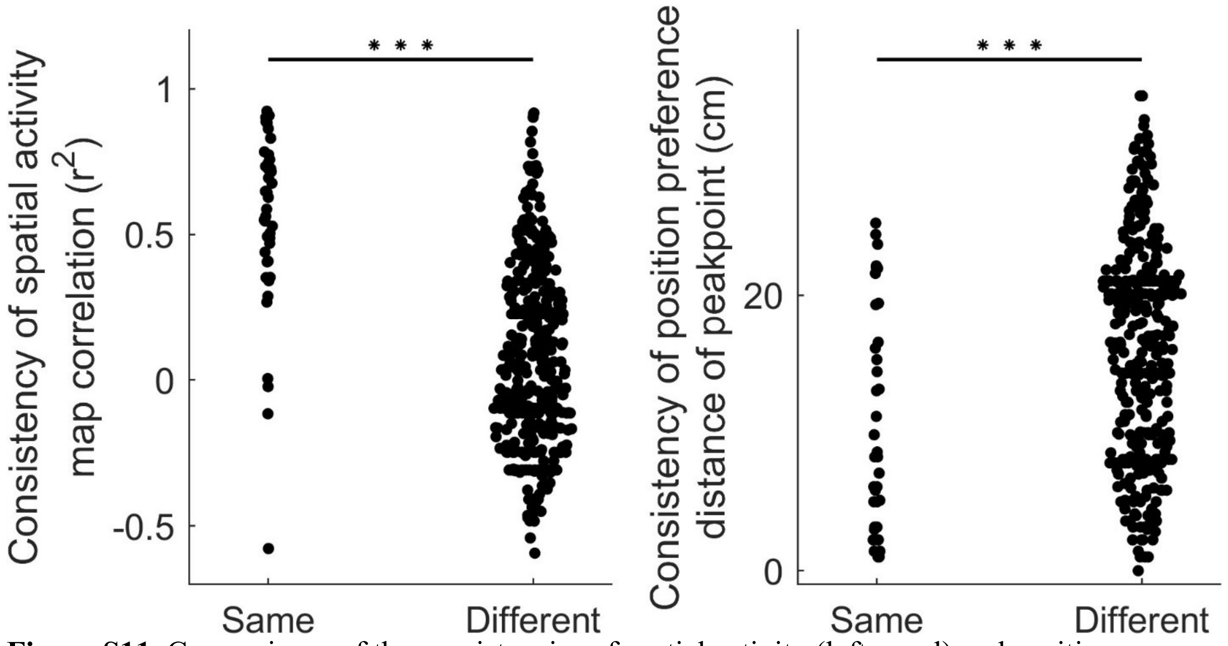


**Figure S11.** Comparisons of the consistencies of spatial activity (left panel) and position preferences (right panel) between the same (*n* = 41 pairs) or different units (*n* = 400 pairs) recorded in different sessions. If a unit was identified being repeatedly recorded across sessions, it had significantly stable spatial firing rate maps and significantly smaller drift in its preferred hand position. *** $P < 0.001$, Kruskal-Wallis test.

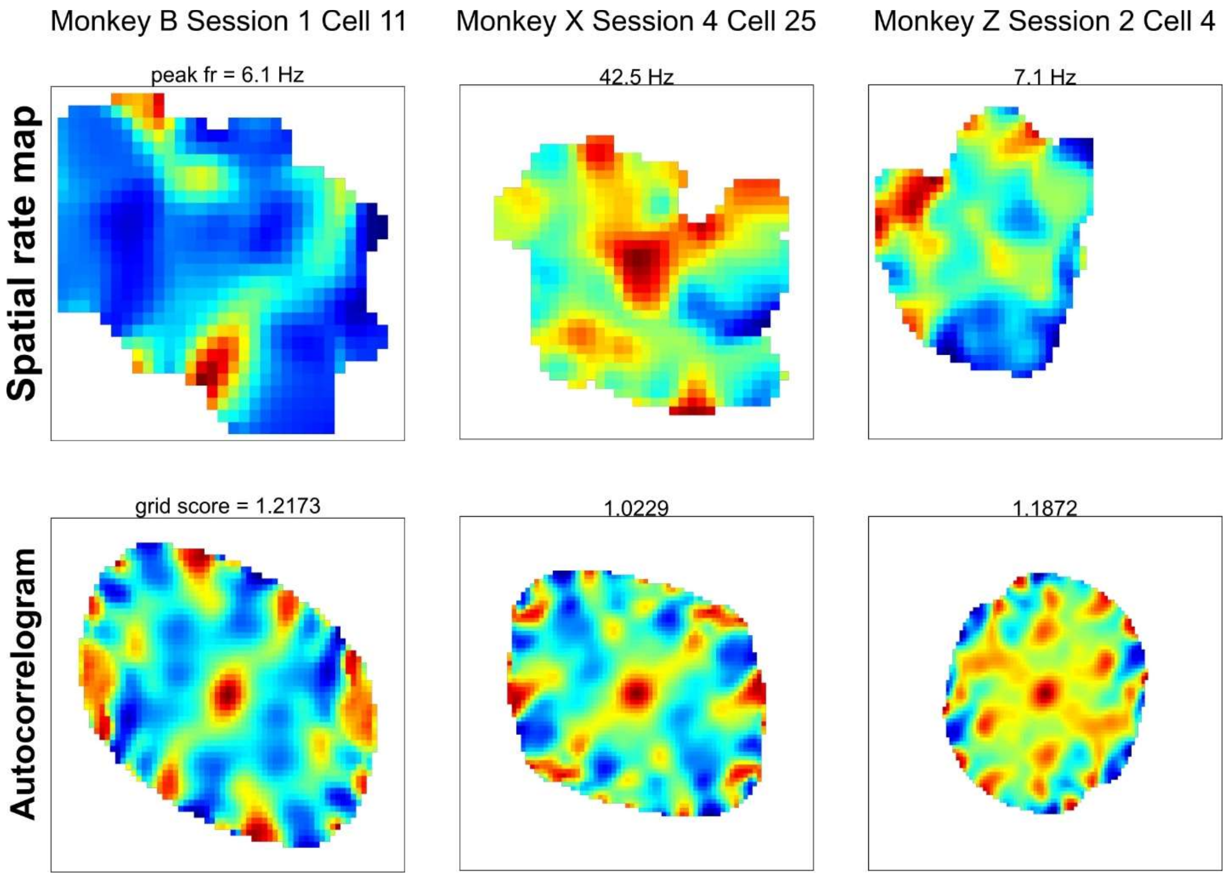


**Figure S12. Grid-like cells in PMd.** 1 grid cell in the XY plane (from Monkey B) and 2 grid cells in the XZ plane (from Monkey X and Z) were identified. The top panels show the 2D firing rate maps with the peak firing rates indicated. The bottom panels show corresponding autocorrelations with the grid scores.
